# PECAn, a pipeline for image processing and statistical analysis of complex mosaic 3D tissues

**DOI:** 10.1101/2021.07.06.451317

**Authors:** Michael E. Baumgartner, Paul F. Langton, Alex Mastrogiannopoulos, Remi Logeay, Eugenia Piddini

## Abstract

Investigating organ biology requires sophisticated methodologies to induce genetically distinct clones within a tissue. Microscopic analysis of such samples produces information-rich 3D images. However, the 3D nature and spatial anisotropy of clones makes sample analysis challenging and slow and limits the amount of information that can be extracted manually. Here we have developed a pipeline for image processing and statistical data analysis which automatically extracts sophisticated parameters from complex multi-genotype 3D images. The pipeline includes data handling, machine-learning-enabled segmentation, multivariant statistical analysis, and graph generation. This enables researchers to run rigorous analyses on images and videos at scale and in a fraction of the time, without requiring programming skills. We demonstrate the power of this pipeline by applying it to the study of Minute cell competition. We find an unappreciated sexual dimorphism in Minute competition and identify, by statistical regression analysis, tissue parameters that model and predict competitive death.

## Introduction

With the advent of safe, non-invasive tools for generating and marking genetically distinct subpopulations of cells within intact organisms, clonal analysis has become a common tool for the study of heterogeneous cell populations and mosaic tissues (Reviewed in ^1^). These tools come in many forms but achieve a common goal: a subset of cells within a tissue, termed a ‘clone,’ acquires a genetic alteration absent in the surrounding tissue. These alterations include mutations in single genes, overexpression of transgenes, and swapping of entire chromosome arms. When these are combined with a visible marker, such as a fluorescent protein, researchers can carry out lineage tracing as well as assess how alterations in a given gene or pathway influence a cell’s behaviour and its interactions with neighbours.

While this technique presents unique avenues for investigating tissue biology, extracting information from mosaic tissues by microscopic analyses is often a bottleneck. The resulting images, typically acquired by 3D confocal microscopy, are information rich and provide insights on phenotypes, such as signal intensity of antibodies or reporters, clone numbers, clone size, shape and apoptosis, all in 3D. Extracting this information, however, is time consuming and complicated. In recent years, a number of powerful image analysis tools have been developed, including new image analysis environments enabling construction of bespoke algorithms, such as CellProfiler^2^, and more targeted software, such as tools for analysis of morphogenesis and cell shape *in situ*^3–5^ and machine-learning enabled segmentation of twin-spot clones^6^, however, there is no image analysis tools designed to analyse complex, multi-genotype images for diverse parameters such as levels of apoptosis, fluorescence and speckle intensity, cell number and density, and parameters of individual clones. Analysis of such confocal images is therefore typically performed in a manual or semi-automated fashion.

Performing such analysis by hand is time-consuming, inconsistent, and error prone, with large datasets often requiring days of work. Researchers must make a compromise between the number of samples they can practicably analyse and the numbers they need to obtain statistically meaningful results. This bottleneck therefore impairs both the speed, quality, and scope of data generation.

Here we present a comprehensive, high-throughput image and data processing pipeline for automating analysis of mosaic experiments: the <u>P</u>ipeline for <u>E</u>nhanced <u>C</u>lonal <u>An</u>alysis (PECAn). This software readily performs myriad clonal analyses, including identifying the number, size, position, and fluorescence properties of clones and single cells by region and genotype in 3D space. In addition, PECAn features an incorporated statistical analysis and graph generation application, allowing users to visualize and evaluate their results in an automated fashion and at scale. This software is flexible, user friendly, and accessible to biologists with no computational or image analysis background, as all inputs are made using graphical user interfaces and require no prior programming knowledge.

To demonstrate the utility of this software, we employed PECAn to analyse parameters of ribosome protein mutant (*Rp/+*) cell competition in *Drosophila* imaginal wing discs – a phenomenon whereby cells carrying heterozygous mutations in ribosomal protein genes, known as *Minutes*, are eliminated from mosaic tissues when proximal to wildtype cells^7^ (Supplementary Figure 1). *Rp/+* cells are therefore said to act as ‘losers’ relative to wildtype ‘winners.’ In the field of cell competition, in particular, there is a pressing need for automated image and single cell analytic techniques^8^. The use of PECAn allowed us to: identify an unappreciated sexual dimorphism in *Rp/+* cell competition, characterize rigorously *Rp/+* cell death properties in competing and non-competing tissues, and identify by logistic regression analysis the tissue parameters that accurately model and predict competitive cell death.

Thus, PECAn reduces operator induced variability and improves speed, consistency, and sensitivity relative to existing techniques while allowing for many new and useful experimental analyses.

## Results

### PECAn design strategy

The PECAn image analysis pipeline consists of two complementary components: (1) a FIJI/imageJ^9^ plugin for analysing images and extracting measurements and (2) an R-Shiny-based web application for processing data, running statistical tests, and generating graphs and plots (Figure 1a). In order to make this software as accessible to biologists as possible, this design prioritizes ease-of-use, requires no prior computational or programming knowledge, and runs entirely on free, publicly available software platforms that are familiar to biologists. All user inputs and processed outputs are made via graphical user interfaces (GUIs) with step-by-step instructions and labels. This program is also optimized for high-throughput batch processing and is capable of analysing large datasets of hundreds of samples with minimal user input. In order to maximize customizability and adaptability, the pipeline incorporates WEKA machine-learning based image analysis^10^. WEKA provides, in FIJI, a GUI-enabled means of training an algorithm using supervised machine-learning, which can classify pixels according to user-defined categories. PECAn links this function directly to image segmentation and measurements. Therefore, images that can be classified by WEKA can be readily incorporated into the pipeline without writing any new code. The pipeline furthermore incorporates tools to allow users to incorporate their own code as modules, therefore custom scripts or additional machine learning segmentation tools can be readily incorporated into the pipeline by anyone with experience in coding using FIJI-compatible languages such as Python and Java.

**Figure 1:**
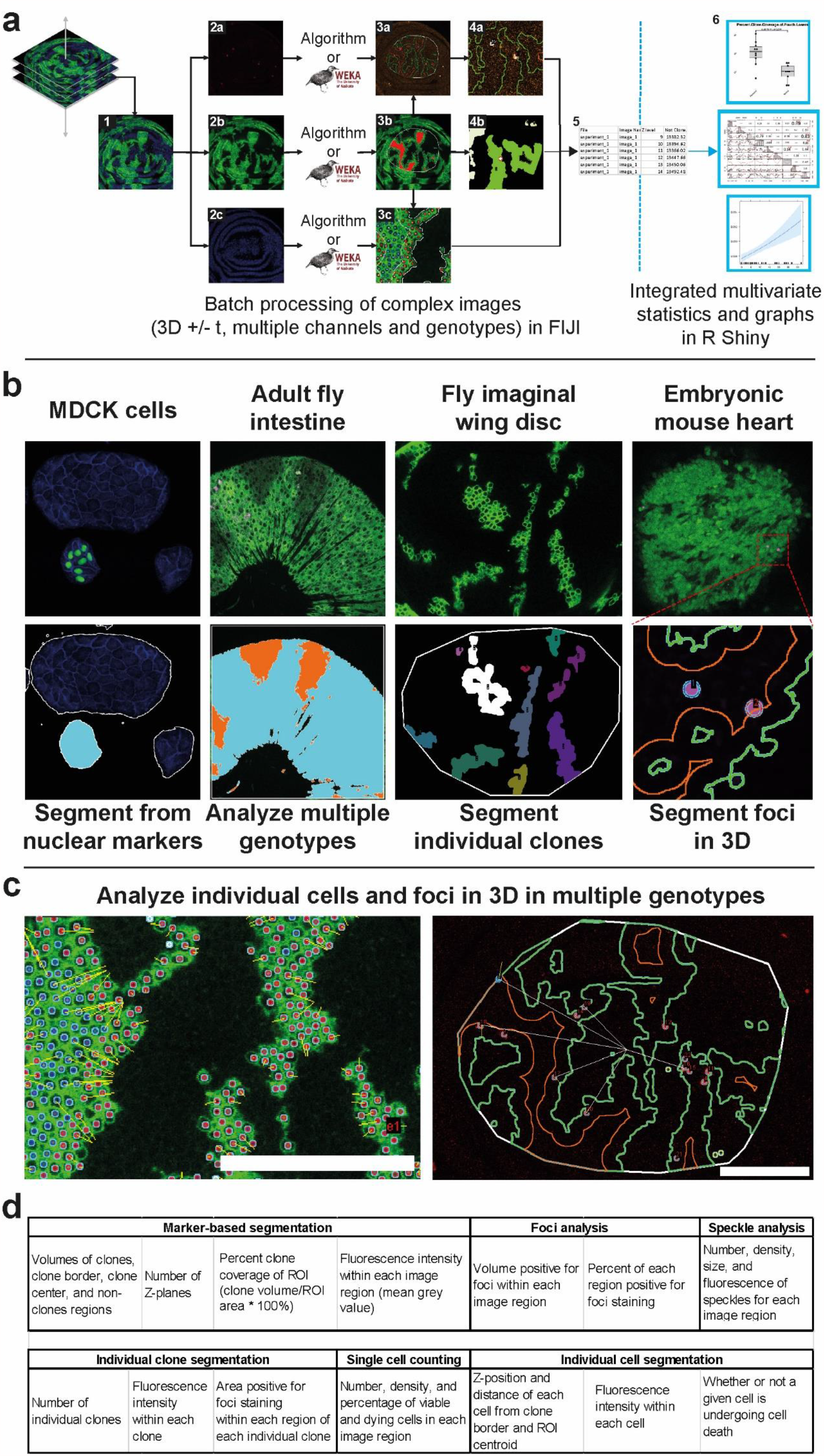
PECAn is a versatile tool for the analysis of complex 3D images. (**a**) Structure of PECAn image analysis software. Images are initially analyzed in FIJI/ImageJ. In order to preserve three dimensional information, each Z-plane (1) of each image is analysed sequentially and compared to adjacent Z-planes. The channels of the image – for instance foci/cell death staining (2a), clones (2b), and nuclear mask (2c) are split apart and processed independently either by the built-in algorithms or by a WEKA classifier. For instance, the foci/cell death mask (3a) can be used to perform 3D segmentation of individual foci and can then be cross referenced against the clone ROIs (3b) to yield a density of foci in different regions within the image (4a) and the level of cell death in individual clones (4b). Furthermore, clone and nuclear stain channels can be used together to identify individual cells in three dimensions (3c). The data are exported as CSV files (5) which can then be uploaded into the R Shiny companion app for statistical analysis and graph generation (6). (**b**) PECAn is able to analyse samples derived from diverse tissues and model organisms. (left) PECAn identifies distinct subpopulations of MDCK cells by combining a nuclear GFP marker with seeded-region growing techniques. (center left) PECAn analyses two distinct genotypes in three dimensions in an adult *Drosophila* intestine sample. (center right) PECAn can segment and analyze individual clones in three dimensions in a *Drosophila* imaginal disc. (right) PECAn identifies density of TUNEL-positive foci in embryonic mouse hearts carrying differently labelled clones. (**c**) PECAn can identify individual cells (left) and individual foci (right) in three dimensions and analyse their position within the tissue, proximity to the clone border (yellow lines), or distance to the clone centroid (white lines). Individual cells/foci are colour coded; a red border denotes a cell in the clone border region, and a blue border denotes a cell in the clone centre region. Scale bars correspond to 50µM. (**d**) Examples of some of the key parameters for which PECAn can assess an image.

### Marker-Based Segmentation

The first step in mosaic analysis is to segment cell populations by visible markers. Since populations are generally marked with a fluorescent signal, identification and separation of regions on the basis of genotype is generally trivial, and the ImageJ/FIJI plugin includes built-in algorithms suitable for two-genotype classification in mosaic tissues. No one algorithm can properly segment all different means by which biologists mark clones. This pipeline therefore integrates WEKA machine-learning enabled segmentation. Thus, PECAn can process any images with clones, which can be segmented either by the built-in algorithm, via WEKA, or by custom user-made code. The pipeline can also process images with a large number of distinctly marked genotypes, making it suitable for assessing clones generated with complex multi-colour genetic tools. To facilitate the analysis of cell interactions, the clonal region is then further subdivided into a peripheral clone border region and the clone centre region.

### Individual Clone Segmentation

The PECAn FIJI plugin allows for analysis of individual clones within an image or tissue (Figure 1b). To do so, the algorithm identifies each disjoint region identified via the marker-based segmentation as distinct ROIs in all Z-planes. The algorithm then compares the ROIs in three dimensions for contiguity while also accounting for instances where clones split apart or merge together across Z-planes. This allows the algorithm to accurately count the size and number of each individual clone, while also assessing them for additional fluorescence-based parameters, such as levels of apoptosis and regional fluorescence intensity.

### Single Cell Segmentation

Segmentation of individual cells operates in a similar fashion. Disjoint regions corresponding to individual cells are segmented either via built-in algorithms or via a WEKA classifier. Each ROI is then compared to ROIs on adjacent Z-planes. Individual ROIs are linked across Z-planes using centroid-based tracking. Individual cells can then be analysed by various metrics, including their position within the tissue, fluorescence intensity, distance to the clone perimeter, and viability as well as by the population-level parameters of the entire tissue (Figure 1c).

### Foci Segmentation

In confocal images, many reporters and stainings present as distinct foci or blobs, rather than as a homogenous signal. Examples of this include assays for cell death and apoptosis, such as antibody stainings for cleaved caspases and TUNEL assays. These methods generally detect dying cells with roughly cell-sized foci, making bulk fluorescence intensity measurements unsuitable. To circumvent this problem, we developed and embedded in PECAn an algorithm for foci segmentation, which uses alternating pixel intensity and size-based filters (Supplementary Figure 2). Alternatively, this can be substituted with a WEKA classifier or custom code for foci detection. It is then possible to calculate Foci enrichment within given regions of the image, such as Regions of Interest (ROI) identified via the marker-based and single clone segmentation steps (Supplementary Figure 2). From this, the algorithm determines the area and percentage of overlap – e.g. the fraction of an ROI that is positive for the cell death reporter, as in (Supplementary Figure 2).

For assays such as apoptosis measurements, however, the traditional metric is to count the number and/or density of discrete foci, to get an estimate of the number of dying cells. This presents additional challenges similar to those faced in tracking individual cells: foci must be accounted for in three dimensions to prevent double or triple counting the same foci, and the algorithm must be able to distinguish between tightly packed foci. This is accomplished using a classic watershed-based approach using the MorphoLibJ toolkit^11^, combined with 3D centroid-based tracking, allowing for precise counts and localization of foci. Individual foci can then be analysed by various parameters, such as their position within the tissue and proximity to various landmarks (Figure 1c). This metric can furthermore be cross-referenced against individual cell counting algorithm to allow the software to determine the exact number of cells positive or negative for a given foci staining in each region of the 3D tissue.

### Fluorescence and speckle measurements

Some fluorescent signals present in images as homogenous stainings or as distinct small speckles. We therefore incorporated accepted fluorescence intensity and speckle analysis functionalities into this pipeline. When combined with the other segmentation modalities present in the software, this allows for measurements of fluorescence intensity and speckle density, number, size and other parameters within different ROIs. Thus, this image analysis pipeline is compatible with a broad range of immunofluorescence and reporter signals, which can be analysed automatically to extract regional properties of the reporters relative to the mosaic composition of the sample. By combining these various modalities, PECAn can rigorously and automatically quantify numerous biologically relevant phenotypes (Figure 1d).

### R-Shiny-Based Web Application

Once the images have been processed, the next bottleneck in data analysis comes in the form of data analysis, statistical testing and plot generation. We therefore developed a companion application in R statistical software to handle the data generated from the FIJI/ImageJ plugin. While R is a powerful statistical software with excellent graphical packages for generating plots and graphs, it is not user-friendly or approachable to someone without prior experience programming in the language. We therefore incorporated our R analysis script into a Shiny web app. This approach combines the statistical power of R with a user-friendly web-based GUI, allowing users to analyse their data without any specialized software or programming experience (Figure 2). This application is designed to automatically process data generated by the FIJI/ImageJ plugin, but can also analyse generic datasets saved as CSV files where data are arranged in columns.

**Figure 2:**
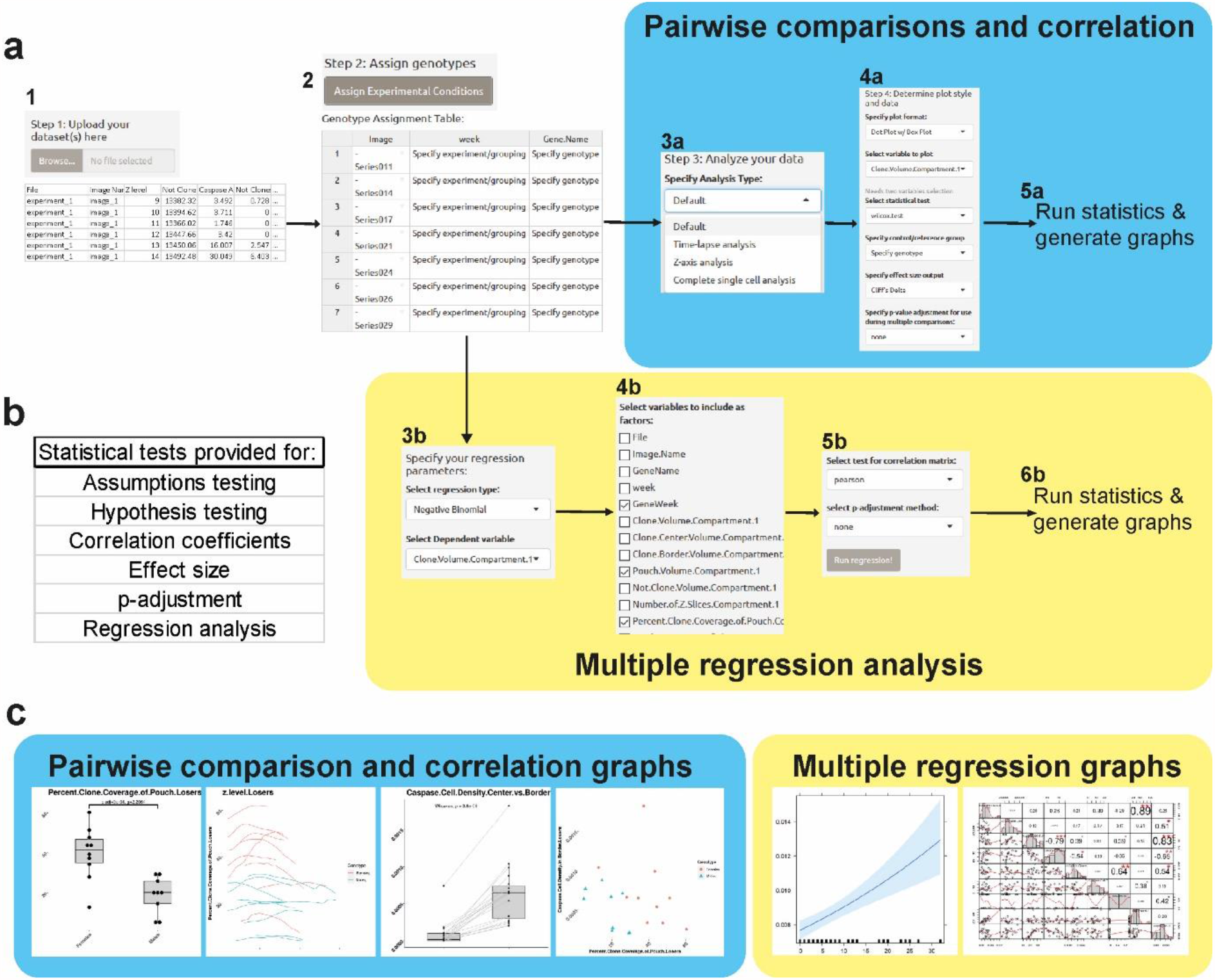
Workflow of the PECAn R Shiny statistical analysis application. (**a**) Users upload the datasets generated by the FIJI/ImageJ plugin (1) and group images by experiment and treatment/genotype/condition using the graphical user interface (2). The user can then assess their samples either via classical uni- and bi-variate tests (a/blue) or via multiple regression analysis (b/yellow). To perform uni- and bi-variate tests, the user specifies an analysis to perform (3a) and then selects an appropriate plot to generate, which tests to run, which reference group to use, and which (if any) p-correction for multiple comparisons to perform (4a). Upon running the analysis (5a), output graphs and statistical tests are performed, along with tests for parametric assumptions. To perform multiple regression analysis, the user specifies which form of regression to run and which parameter should act as the dependent variable (3b). The user then selects which parameters to include as predictor variables (4b), which test to use for generating a correlation matrix, and which (if any) p-correction for multiple comparisons to perform (5b). Upon running the analysis (6b), the regression is run along with test for assumptions and both output and diagnostic plots are generated. (**b**) A summary of the kinds of tests supported by the app. (**c**) Examples of plots generated by the app, using the ggplot2 package in R. Different plots are generated for uni- and bi-variate tests (left/blue) and for multiple regression analysis (right/yellow).

Once the data generated from the FIJI/imageJ plugin is uploaded and the user specifies the genotypes, experimental groupings, and the desired analysis, the app automatically runs appropriate statistical tests and generates output plots using a GUI accessible on any conventional web browser. The user is prompted to upload the data files generated by the FIJI plugin, at which point the user groups the data according to condition and/or experiment and specifies how to analyse the data. Once the data is processed, the user is free to assess the dataset in two general ways: (1) classical uni- and bi-variate analysis or (2) multiple regression.

For uni- and bi-variate analysis, the user specifies which variety of graph to generate, which variable(s) to analyse, which statistical test to run (such as a t-test, ANOVA, or correlation coefficient), which effect size metric to run, and if/how to adjust p-values for multiple comparisons. Once specified, the app automatically runs all specified tests along with parametric assumptions tests and generates appropriate graphs. These graphs are customizable and can be exported as publication quality images using the ggplot2 package in R. For multiple regression analysis, the user selects between various techniques, such as logistic, linear, and Poisson regression, and specifies which variables to act as dependent and predictor variables. Upon running the analysis, the app automatically runs appropriate assumptions tests, produces effects plots, and generates a suite of diagnostic plots and metrics to enable the user to evaluate the quality of their analysis. The app furthermore provides tools for performing data transformations. Should the user desire to run any statistical tests not supported by the app, the processed data can be exported and therefore readily incorporated into any conventional statistical analysis software.

In order to ensure that the multiple regression functionalities in the R Shiny analysis app were functioning properly, we ran publicly available tutorial datasets for multiple logistic regression, Poisson regression, negative binomial regression, and linear regression through the analysis app. These analyses generated the expected results (Supplementary Table 1), indicating that the statistical packages had been properly incorporated into the application.

### PECAn Fiji Validation

Upon completion of the pipeline, we sought to validate PECAn by challenging it in three separate ways: (1) using semi-synthetic images, (2) comparing with human analysis, and (3) testing its sensitivity at identifying known strong or mild modulators of cell competition. For these comparisons, we used as reference data sets 3D, multi-channel confocal images of the pouch region of *Drosophila* imaginal wing discs, a roughly planar pseudo-stratified epithelium. In these wing discs, mutant cells carrying heterozygous mutations in the ribosomal small subunit protein *RpS3* gene are generated in a mosaic fashion alongside wildtype cells. We used a flp-out based construct, which generates <u>M</u>inute clones in a <u>w</u>ildtype <u>o</u>rganism (MiWO) (Supplementary Figure 3a and ^12^). In MiWO wing discs, *RpS3*^*+/-*^ cells then undergo Minute competition^12^.

Semi-synthetic images provide a means of testing a large program for errors in the code and data output. In short, images with known properties are fed into the pipeline, and the outputs are compared against the known input values (Figure 3a). The pipeline yielded results consistent with the input values across all parameters tested (Figure 3b), indicating that the software successfully computes the right operations and produces the desired measurements.

**Figure 3:**
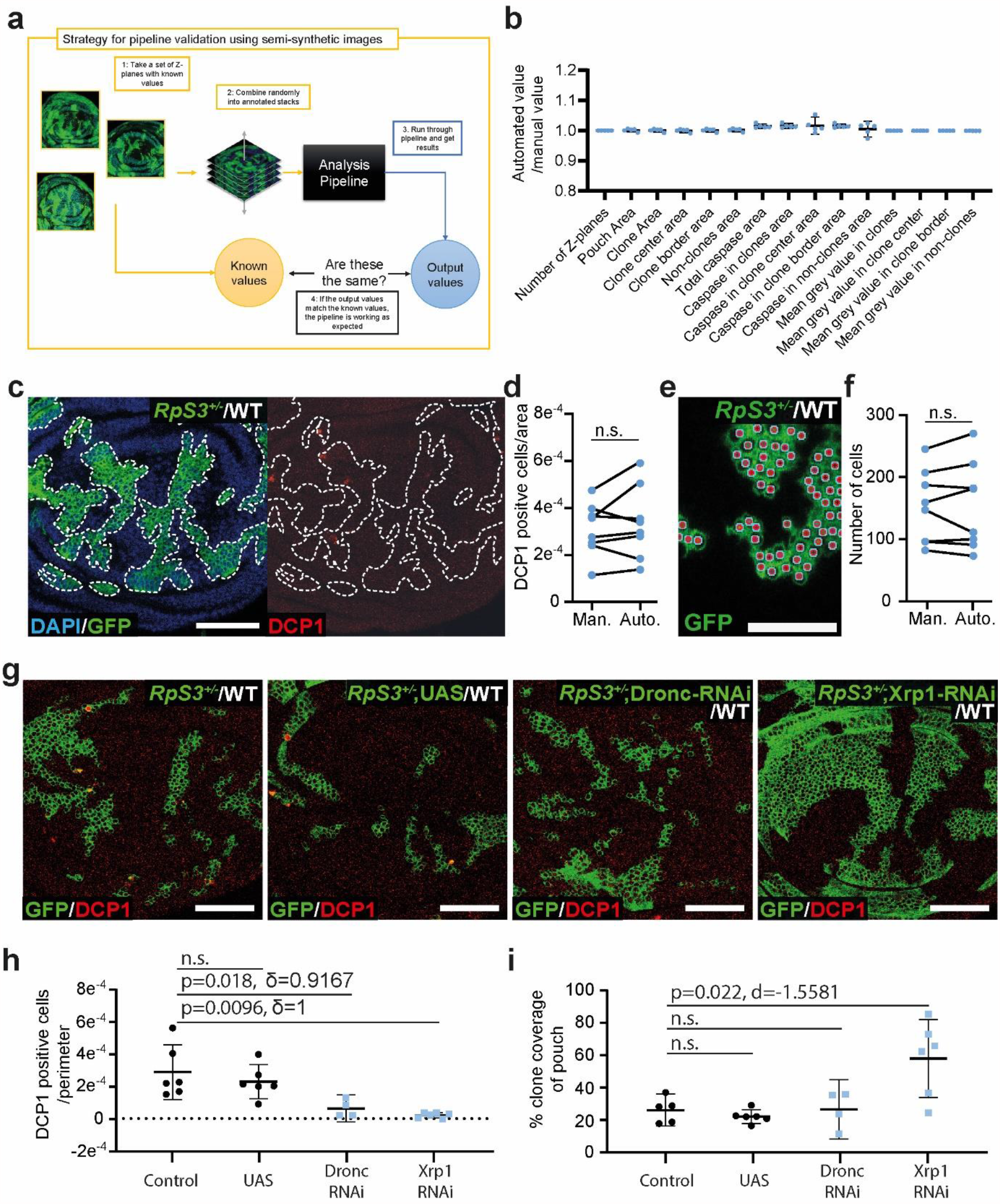
Validation of the PECAn pipeline. (**a**) Schematic of the semi-synthetic image validation strategy. Z-planes with known parameters were randomly combined into Z-stacks of a random length. These known images were analysed using the macro, and the macro results were compared against the known values. This approach helps detect errors in the measurements and outputs of the macro. (**b**) Results generated by the macro divided by the known value. All values closely match their expected values, with deviations attributable to rounding errors. (**c**) Representative image of a competing wing disc stained with anti-cleaved DCP1. (Left) *RpS3*^*+/-*^ loser clones are marked in green, and DAPI staining in blue. (Right) The anti-cleaved DCP1 staining is shown in red and the dotted line represents the outline of the loser clones. (**d**) Comparison of the density of apoptotic events in the clone border as determined by hand against the macro outputs. (**e**) Sample output of individual cell counts performed by PECAn in a MiWO wing disc. Individual cells are marked with a uniquely color-coded stamp. (**f**) Comparison of quantifications generated manually against those generated by the macro. (**g**) Representative images of competing *RpS3*^*+/-*^ cells (green) with a stain for cleaved-DCP1 (red). From left to right, the genotypes are a control expressing no transgenes other than GFP, a negative control driving expression of an empty UAS promoter, a positive control expressing an RNAi against Dronc, an initiator caspase, and a positive control expressing an RNAi against Xrp1. (**h**) Automated quantification of the density of cleaved-DCP1-positive cells in the clone border region. (**i**) Automated quantification of the percentage of the wing disc pouch occupied by *RpS3*^*+/-*^ cells. Scale bars correspond to 50µm.

To compare the pipeline’s performance against human analysis, an image dataset was both analysed by hand and run independently through PECAn. The results generated by the macro were consistent with those generated by users: both the density of apoptosis in the clone border region (Figure 3c-d, Supplementary Figure 3b) and the counts of individual cells (Figure 3e-f) were not significantly different, showing that the pipeline performs comparably to manual quantification.

The pipeline was then tested for its ability to identify known modulators of Minute cell competition. *Dronc* and *Xrp1* knockdown have both been shown to inhibit Minute cell competition: *Dronc* RNAi yields a mild rescue, as it only inhibits *Rp/+* cells from competition-induced apoptosis^13^,whereas mutation in *Xrp1* yields a strong rescue, as it inhibits both competitive death and the slow growth phenotype of *Rp/+* cells^14,15^. Thus, for this test, we generated control MiWO clones and MiWO clones expressing RNAi against those cell competition modulators. These samples were fed into the pipeline and assessed for density of apoptosis at the clone border and for clone size, the two common metrics used to assess Minute cell competition. The pipeline successfully detected both known modulators and distinguished these from controls. Importantly, PECAn distinguished between the rescue modalities of these two genetic manipulations, scoring a rescue in both clone size and competitive death for Xrp1 and a rescue in death but not in clone size for *Dronc* RNAi. This shows that this analysis pipeline detects automatically both strong and mild modulators of Minute cell competition (Figure 3g-i).

### PECAn reveals sexual dimorphism in Minute competition

Having successfully validated PECAn, we then sought to utilize this new toolset to investigate at unprecedented scale and sensitivity some of the parameters of Minute cell competition. Sexually dimorphic phenotypes are common in *Drosophila* cell biology^16,17^, but this has not been investigated for cell competition. To ask whether Minute cell competition displays sexual dimorphism, we dissected separately male and female larvae from (1) a cross of wildtype males bred with females carrying the MiWO transgenes, and (2) a cross of wildtype females bred with males carrying the MiWO transgenes (Figure 4a-c). No differences were observed between female larvae, regardless of parental genotype. Surprisingly, however, we observed a strong sexual dimorphism: *RpS3*^*+/-*^ clones covered a smaller proportion of the tissue in males relative to females (Figure 4a-b). Interestingly, this effect was confined to clone coverage, as we did not observe differences in the frequency of competition-induced apoptosis between males and females (Figure 4a,c). Thus, cell competition in this system is highly sexually dimorphic.

**Figure 4:**
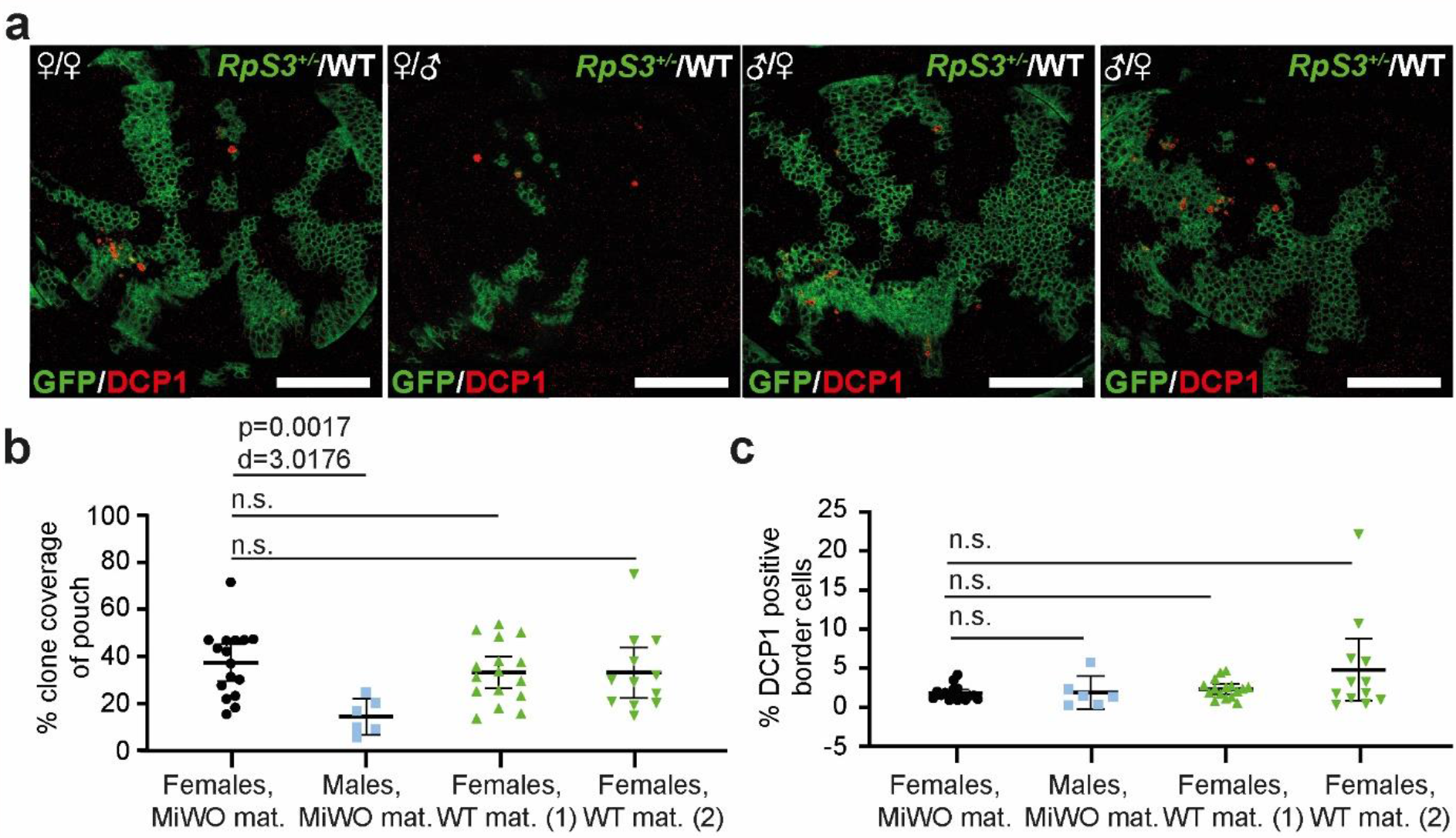
Competing RpS3+/- losers exhibit a further growth disadvantage in male wing discs. (**a**) Representative images of MiWO wing discs containing *RpS3*^+/-^ losers (green) competing against wildtype winners (unlabeled) and stained for cleaved-DCP1 (red). Wing discs were derived from female (left) or male (center left) larvae who inherited the MiWO construct from their mothers, or from two separate dissections of female larvae who inherited the MiWO construct from their fathers (center right and right). (**b**) Quantification of loser clone coverage in wing discs as in (**a**). (**c**) Quantification of the percentage of cells undergoing apoptosis at the loser clone border in wing discs as in (**a**). Scale bars correspond to 50µm.

### Quantitative analysis of the parameters of Minute competitive apoptosis using PECAn

Next, we exploited the sensitivity and high-throughput analysis capabilities afforded by PECAn to study the properties of competition-induced cell death in *Rp/+* cells. It is well established that competing *Rp/+* cells proximal to wildtype winners exhibit higher levels of cell death relative to distal competing *Rp/+* cells^12,13,18,19^. It is also well established that non-mosaic *Rp/+* cells exhibit elevated levels of cell-autonomous apoptosis^12,20–22^. However, it is not known how these levels of cell-autonomous apoptosis compare to those in competing *Rp/+* cells. We therefore generated wing discs with a mosaic anterior compartment and an entirely *Rp/+* posterior compartment. As Minute cell competition does not occur across compartment boundaries^23^, this allowed us to compare competitive and non-competitive cell death within a single tissue (Figure 5a). We then analyzed these samples in PECAn, utilizing its ability to assess multiple ROIs within a single image (Figure 5b). The density of apoptotic cells was several folds higher at the competing *Rp/+* clone border relative to the internal reference death level observed in non-competing posterior compartment, indicating that competitive cell death represents an elevation over the non-competing condition (Figure 5c).

**Figure 5:**
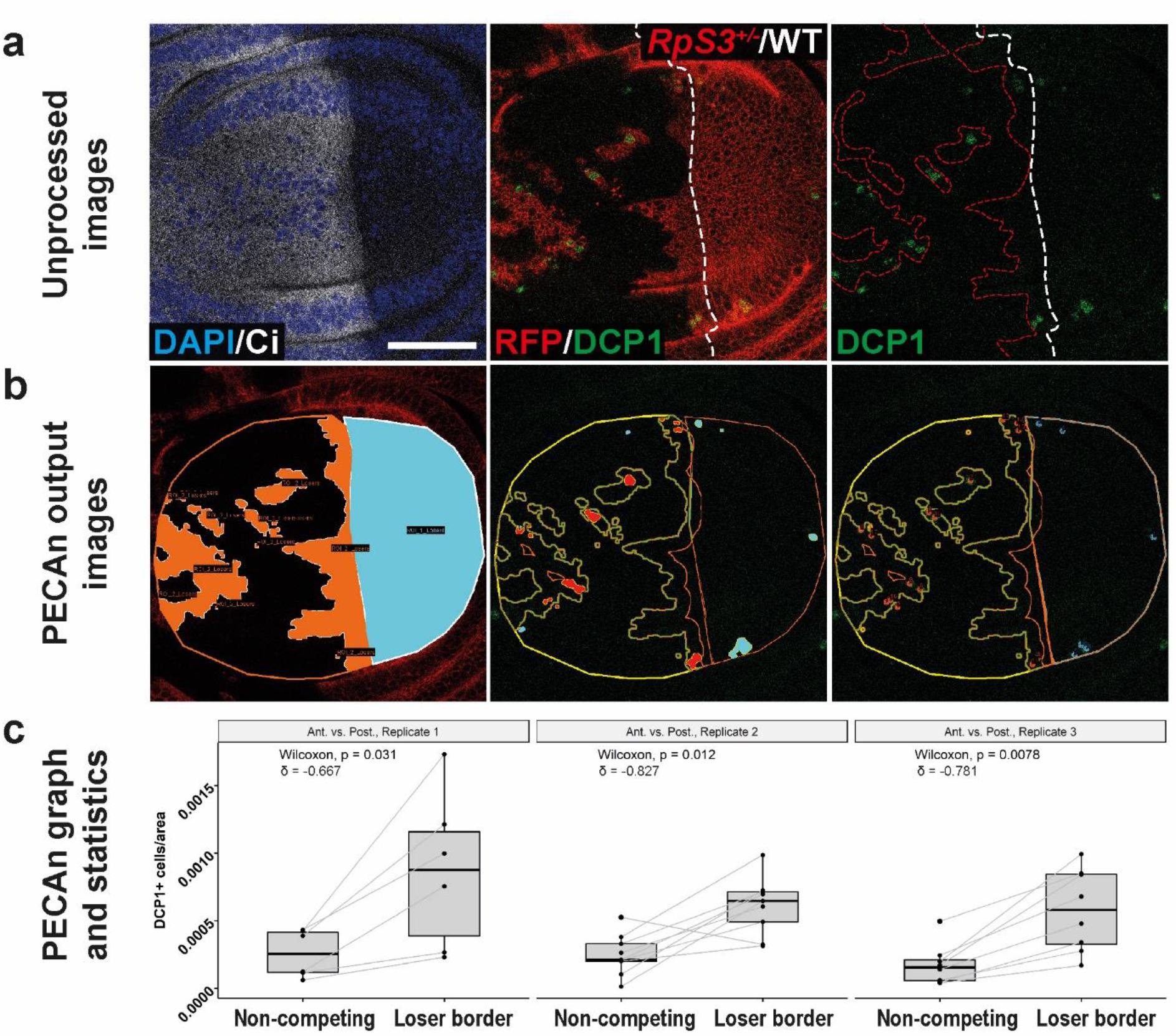
*RpS3*^*+/-*^ cells exhibit higher levels of cell death during competition than in a non-mosaic context. (**a**) Wing discs carrying a mosaic anterior compartment (identified by the anterior fate marker, Ci, shown in white) containing both *RpS3*^*+/-*^ (red) and wildtype (unlabelled) cells and a posterior compartment (negative for Ci) that is entirely *RpS3*^*+/-*.^ These samples were assessed for cell death via a staining for cleaved-DCP1 (green) and analyzed using PECAn. (**b**) PECAn output images identifying *Minute* cells in the anterior (orange) and posterior (cyan) compartments (left), the regions of the image positive for the DCP1 staining (middle) and the counts of individual DCP1-positive cells (right). (**c**) Output graphs and statistical tests generated by PECAn showing the density of DCP1-positive cells in the non-competing posterior compartment as compared to the loser clone border in the anterior compartment for three separate experimental replicates. The scale bars correspond to 50µm.

We then sought to use PECAn to rigorously identify the parameters of cell death within wing discs undergoing *Rp/+* competition. We compiled a large dataset of competing MiWO wing discs prepared using consistent conditions, but dissected, processed, and imaged in separate batches (67 wing discs and 192,207 *RpS3*^*+/-*^ cells from seven separate dissections). To control for the observed sexual dimorphism, only female larvae were dissected. Individual cells within all wing discs were assessed for their cell death (using PECAn single cell and foci segmentation) and spatial properties, and the resulting dataset was subjected to a multiple logistic regression-based analysis, using the PECAn statistical analysis app, with the dependent variable being whether or not a given cell is viable or apoptotic (Figure 6a-f, Supplementary Table 2).

**Figure 6:**
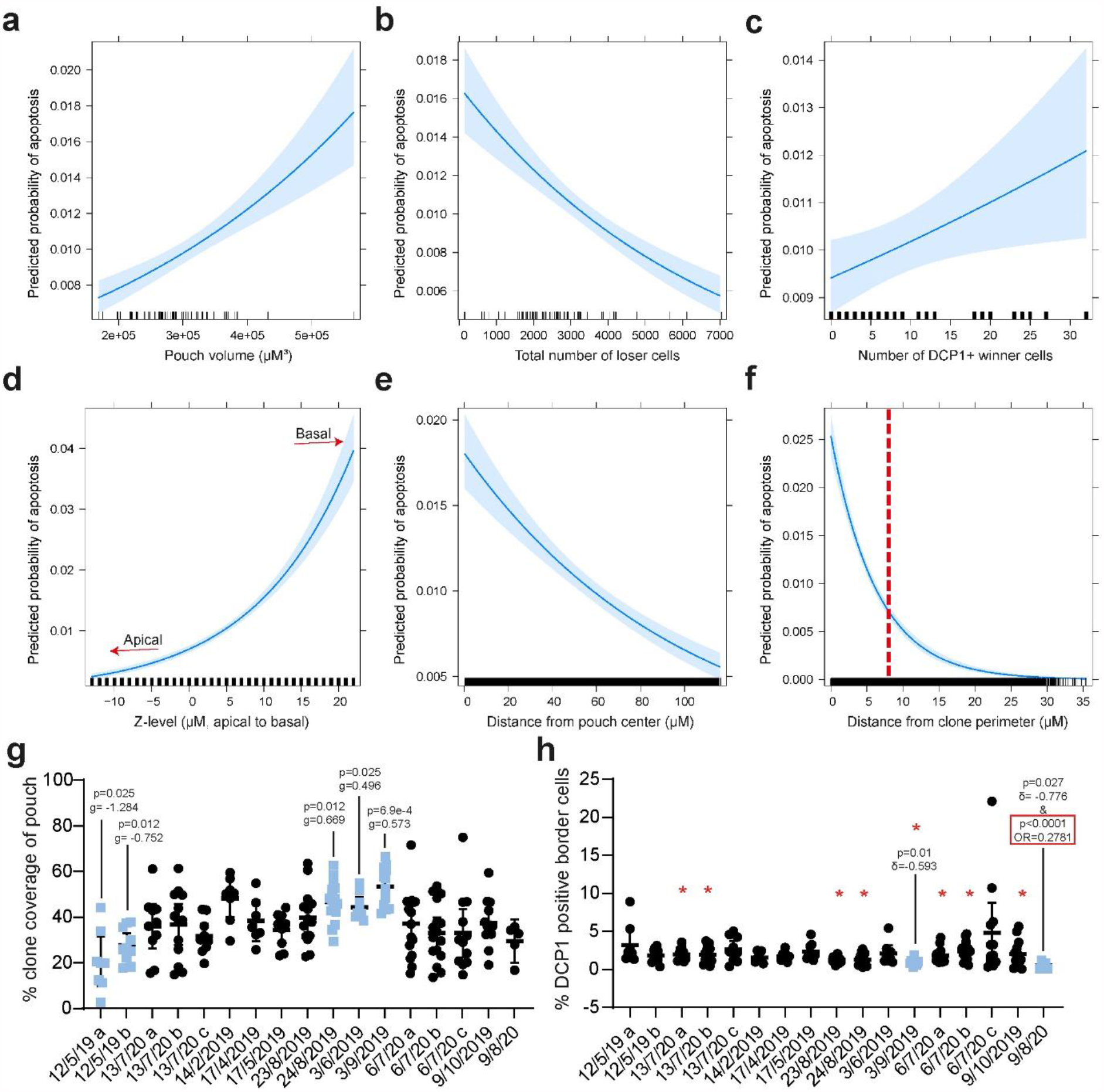
Multiple logistic regression of competing wing discs reveals tissue parameters influencing loser cell death. (a-f) Predicted effects plots of logistic regression analysis of individual cells in competing wing discs shown in Supplementary Table 2. In these plots the probability of cell death is plotted against the volume of the pouch (**a**), the total number of *RpS3*^*+/-*^ loser cells in the sample (**b**), the number of DCP1-positive wildtype winner cells in the sample (**c**), the Z-level of the cell (**d**), the distance of the cell from the centre of the pouch (**e**), and the distance of the cell from the loser clone perimeter (**f**). These plots show the expected probability of apoptosis upon changing a given predictor variable – the distance of the cell from the clone perimeter, whilst all other predictors are set to their mean value. Pale blue region denotes the 95% confidence interval, and black tick marks on the x-axis denote observed values in the dataset. The red dashed line in (**f**) is at 8µm, which corresponds roughly to 2 cell diameters from the clone border and denoting the clone border/clone centre cut-off used in other analyses in this study. (**g**-**h**) Comparison of replicates/experiments used in the logistic regression analysis shown in Supplementary Table 3. (**g**) The percent loser clone coverage of the wing disc pouch region for each wing disc is shown on the y-axis, while the x-axis shows the replicate/experiment to which they belong, labelled according to the date at which they were imaged. p-values and g-values reflect two-sided student’s t-tests with FDR correction and Hedges’ g effect size metrics relative to the basemean. (**h**) The percentage of DCP1-positive loser cells is shown for each replicate/experiment. Statistics reflect either a two-sided Mann-Whitney U-test with FDR correction along with Cliff’s δ effect size metric, relative to the basemean (no border) or the logistic regression analysis along with odds ratio effect size metrics (OR) shown in Supplementary Data Table 2 (red border). Datasets that were outliers by Mann-Whitney U-test are shown in blue, and red asterisks reflect datasets that had significant p-values by logistic regression but did not meet the effect size cutoff.

This analysis provides evidence for several interesting interactions within competing wing discs. We found that the probability of *Rp/+* cell death varies with the size of the wing pouch region (Figure 6a, Supplementary Table 2). This could suggest that competitive death is more pronounced in slightly older discs. We also found that the rate of loser cell death declines as the number of *RpS3*^*+/-*^ cells increases (Figure 6b, Supplementary Table 2). As this analysis accounts for physical distance of *RpS3*^*+/-*^ cells to the winner, this reduction in death is not exclusively due to a relative decrease in the exposure of loser cells to winners and could suggest that the strength of competition is sensitive to cell community effects influenced by the relative abundance of winners and losers, as it has been reported for Rab5 cell competition^24^. Alternatively, the anticorrelation of loser cell death and loser cell abundance could simply reflect an underlying variability in the intensity of cell competition across wing discs, whereby weaker cell competition would both result in bigger clones and in less competitive death.

Loser cell apoptosis also positively correlates with levels of apoptosis seen in winner cells (Figure 6c, Supplementary Table 2), indicating that loser cells are more likely to undergo apoptosis in wing discs with higher overall levels of cell death. The probability of loser cell death further depends on a cell’s position within the tissue, with basal locations associated with a higher probability of apoptosis (Figure 6d, Supplementary Table 2). This result is consistent with established dynamics of cell death in wing discs generally and during *Rp/+* competition, specifically^18,25,26^.The probability of observing apoptosis also declines the further a loser cell is from the centre of the pouch (Figure 6e, Supplementary Table 2). This observation is consistent with the observation that Minute clones are more aggressively eliminated from the centre of the pouch^27^.

Lastly, this analysis also identifies that loser cell death correlates strongly with physical proximity to winner cells. As multiple logistic regression analysis accounts for possible confounding factors in the dataset (e.g. clone size, shape, position, volume etc), it is significant that this analysis, while considering the relative contributions of these factors, finds the probability of loser cell death increases exponentially the closer a cell is to the wildtype (Figure 6f, Supplementary Table 2). This relationship is consistent with previous analyses, which found that rates of loser cell death were highest within one-to-two cell diameters of the clone border^13,18^. These results confirm that the border death seen in competing wing discs is a fundamental feature of *Rp/+* cell competition.

### PECAn statistical regression analysis faithfully models and predicts competitive cell death

While this logistic regression model of loser cell apoptosis provides results consistent with the experimental literature, the model has a weak point which must be addressed: a middling Nagelkerke pseudo-R^2^ value of 0.257. In the context of this particular model, the pseudo-R^2^ can be thought of as a measure of how much better the model is at predicting which cells are apoptosing than a null model. This model, therefore, is poor at determining which cells will be apoptotic. There are two possible explanations for this observation: either there is a key predictor variable missing from this model, or the dependent variable has a high level of stochasticity. As the probability that a given cell will be apoptotic is low (1.4% of cells in the dataset are apoptotic), it seems likely that there is a strong element of stochasticity to exactly which cells are apoptotic at a given moment in time and that the model would struggle to accurately predict a rare, stochastic event. If the latter hypothesis is correct then the model should accurately predict the frequency of apoptotic cells at the level of the entire wing disc, as a population analysis would dilute the impact of stochasticity.

We therefore conducted a logistic regression analysis on the number of cells that are undergoing apoptosis versus the number that are non-apoptotic at MIWO competing clone borders (Supplementary Table 3). The set of predictor variables used for this analysis was updated as shown in Supplementary Table 3, as some of the parameters used in the prior model were unsuited to this analysis. As this analysis was less computationally expensive, we could use an expanded dataset corresponding to 183 wing discs from 17 separate dissections. The resulting analysis is able to predict the number of apoptotic and non-apoptotic loser cells at clone borders in each wing disc with high accuracy, with a Nagelkerke pseudo-R^2^ value of 0.9988 – indicative of an excellent fit between the model and the data. These results together indicate that, while rates of loser cell apoptosis increase according to several predictor variables, precisely which cells will undergo apoptosis is a stochastic process. The goodness-of-fit between this model and the data further indicates that this set of predictor variables provides a robust and comprehensive assessment of the parameters dictating levels of loser cell death at the clone border.

With this second analysis, we were furthermore in a position to evaluate how consistent metrics of cell competition are across experimental replicates, using both classical univariate and multiple logistic regression based techniques (Figure 6g-h, Supplementary Table 3). This furthermore provides an important control for the logistic regression analyses here presented: as these analyses were based on an assessment of cell death across separate samples and replicates, the quality of the conclusions derived can only be as good as the quality of the dataset. Any numbers derived from an inconsistent dataset would risk overfitting noise. These replicates show considerable variability in the proportion of the pouch covered by *RpS3*^*+/-*^ (Figure 6g). This is to be expected, as loser clones in this system are induced in a semi-stochastic fashion and these experiments are not rigorously time controlled. These replicates, however, exhibit a high level of consistency at the level of border cell death, with only two of seventeen datasets – 3/9/19 and 9/8/20 - showing significant deviation from the basemean, as determined via a Mann-Whitney U test with an FDR p-adjustment and Cliff’s δ effect size comparison (Figure 6h). When assessed by multiple logistic regression, many replicates yielded significant p-values, though only one replicate had a non-negligible odds ratio effect size^28^ (Figure 6h). These data indicate that competitive loser cell death in *Rp/+* competition is a robust metric of competition that is refractory to noise across replicates.

## Discussion

Clonal analysis techniques *in situ* provide unique and powerful methods for probing the behaviour of and interactions between heterogeneous cell populations. The challenges of extracting meaningful data from these samples, however, limits the speed, consistency, and quality with which researchers can conduct these experiments. The time consuming and low-resolution nature of extant analytical techniques also constrains the number and type of analyses that can be conducted, leading to under-analysed datasets, operator dependent variability/reproducibility issues, and low statistical power. To address this, we have created a complete high-throughput image and data analysis pipeline for the analyses of mosaic samples. This pathway is approachable and user-friendly even to biologists with no programming or image analysis expertise.

On installation, the software can readily assess clones for multiple parameters such as cell death, fluorescence intensity, and speckle density in three-dimensions. It can furthermore perform these analyses on a range of experimental systems. The range and utility of this system is further expanded by the incorporation of WEKA image segmentation, allowing for broad applicability of this system to images of tissue samples and fixed cells.

A key strength of this software is that it is streamlined and integrated, while still retaining flexibility and customizability. The underlying code is written in the powerful yet approachable Jython/Python and R languages, and researchers with programming experience will be able to readily incorporate new tools and algorithms into the pipeline. As the image analysis software is integrated into the powerful FIJI/imageJ software, users without computational backgrounds can expand the functionality of this pipeline by pre-processing samples as necessary. The software furthermore generates clear visual outputs of all measurements performed allowing users to easily and confidently tailor the analyses to their samples.

The identification of a previously undescribed sexual dimorphism in *Rp/+* cell competition also highlights the utility of a high-throughput analysis approach. A more rigorous investigation of this phenomenon is warranted, as the field generally has not accounted for gender before. It is worth noting that this result may be due to differences in clone generation resulting from differing expression of the heat-shock-inducible flippase, as this construct is carried on the X chromosome in our system. This one factor might contribute to noise/variability seen in prior competition experiments. In addition, it may hamper sensitivity and discovery power. This is particularly relevant in screens for novel cell competition genes, where clone size is normally the parameter of choice^29–31^, given that more labour-intensive analyses, such as for competitive death cannot be carried out at scale, in the absence of an automated analysis tool like the one presented here.

It is furthermore worth emphasizing that, in this study, conditions were identified which influenced *Rp/+* loser cell death without affecting *Rp/+* loser cell clone size (Dronc RNAi), and vice versa (larval gender). This confirms prior work which shows that influencing one trait need not necessarily influence the other and highlights the fact that *Rp/+* loser cell death and growth are distinct arms of the competition process.

A further advantage of this pipeline is that, with its built-in cell segmentation tools, it enables direct counts of apoptosing and non-apoptosing cells. Such an approach enables the use of more powerful and informative statistical techniques and analyses, such as those presented in this manuscript. These analyses have enabled us to investigate the parameters of competitive cell death with an unprecedented level of quantitative rigor. PECAn, thus, enables researchers to conduct rigorous characterizations of biological phenotypes and assess the potential impact of myriad biologically relevant parameters therein.

## Acknowledgements

We wish to thank Rafael Carazo-Salas and his laboratory for their invaluable guidance in designing, building, and validating the image analysis pipeline. We also thank Stephen Cross and Anatole Chessel for their help in evaluating the software. We also wish to express our gratitude to Daniel Lawson and Susan Connolly for assisting us in implementing, interpreting, and understanding statistical packages in R. Thanks to Cristina Villa del Campo and Miguel Torres for sharing sample images of a postnatal mouse heart. Finally, we thank the Wolfson Bioimaging Facility for access to microscopes. This work was supported a Cancer Research UK Programme Foundation Award to E.P. (Grant C38607/A26831) and a Wellcome Trust Senior Research Fellowship to E.P. (205010/Z/, 16/Z).

## Materials and Methods

### Software development

The Pipeline for Enhanced Competition Analysis (PECAn) software was designed in the script editor of ImageJ299 FIJI version 1.53d^9^. All code was written in the Jython programming language. In addition to base FIJI packages, the Bio-Voxxel, MorphoLibJ^11^, and the IJ-plugins toolkit were used. The R shiny analysis application was made in RStudio using R3.6.3 and shiny 1.5.0. The analysis app uses the following external libraries: PerformanceAnalytics, ggplot2, boot, MASS, car, RColorBrewer, ggpubr, markdown, ggsignif, rhandsontable, msm, dplyr, magrittr, ICSNP, mvnormtest, psych, corrplot, rcompanion, stringr, effsize, sandwich, ggthemes, shinyBS, reshape2, and effects. A list of statistical tests along with their associated R function is provided in the table below. The software, source code, sample images, and instructional videos are provided in the supplementary electronic materials.

| Test | Function | Package |
| --- | --- | --- |
| Wilcoxon Signed Rank | compare_means | ggpubr v0.4.0 |
| Wilcoxon Rank Sum | compare_means | ggpubr v0.4.0 |
| Student's t-test | compare_means | ggpubr v0.4.0 |
| Paired t-test | compare_means | ggpubr v0.4.0 |
| Kruskal-Wallis | compare_means | ggpubr v0.4.0 |
| ANOVA | compare_means | ggpubr v0.4.0 |
| Shapiro-Wilks | shapiro.test | stats v3.6.2 |
| Fligner-Killeen | fligner.test | stats v3.6.2 |
| Fisher's exact | fisher.test | stats v3.6.2 |
| Cliff's delta | fliff.delta | effsize v0.8.1 |
| Cohen's d | cohen.d | effsize v0.8.1 |
| Hedge's | cohen d | effsize v0.8.1 |
| Spearman's rho | cor.test | stats v3.6.2 |
| Pearson's r | cor.test | stats v3.6.2 |
| Kendall's Tau | cor.test | stats v3.6.2 |
| P-adjustments | p.adjust | stats v3.6.2 |
| Correlation matrix | chart.Correlation | PerformanceAnalytics<br>v2.0.4 |
| Multiple linear regression | Lm | stats v3.6.2 |
| Logistic regression | Glm | stats v3.6.2 |
| Poisson regression | Glm | stats v3.6.2 |
| Negative binomial regression | glm.nb | MASS |
| Variance inflation factors | Vif | car v3.0-10 |
| Durbin-Watson Test | durbinWatsonTest | car v3.0-10 |
| Non-constant variance of error | ncvTest | car v3.0-10 |
| Nagelkerke pseudo-R<sup>2</sup> | Nagelkerke | rcompanion v2.3.26 |
| Cox and Snell pseudo-R<sup>2</sup> | Nagelkerke | rcompanion v2.3.26 |
| McFadden pseudo-R<sup>2</sup> | Nagelkerke | rcompanion v2.3.26 |

### Fly husbandry and clone induction

All flies were reared on wheat-based food prepared according to the following recipe: 7.5g/L agar powder, 50g/L baker’s yeast, 55g/L glucose, 35g/L wheat flour, 2.5% nipagin, 0.4% propionic acid and 1.0% penicillin/streptomycin, and all experimental crosses were kept in an incubator at 25°C. The parent cross was allowed to lay eggs for roughly 24 hours, and all dissections were performed on larvae at the wandering third instar stage six days after egg laying. Clone induction was accomplished using a heat-shock inducible FLP recombinase by placing the fly vials in a water bath set to 37°C. MiWO flies were heat shocked for 25 minutes on day 3 after egg laying and mitotic clones were heat shocked for 15 minutes on day 3 after egg laying. For clones expressing temperature-sensitive constructs, fly vials were transferred to a water bath set to the specified temperature immediately following clone induction and were then dissected as normal. Experimental genotypes, and clone induction conditions are listed in Supplementary Table 4.

The following fly stocks were used: RpS3[Plac92] (cat#BL5627, Bloomington Drosophila Stock Center), UAS-Xrp1-RNAi (cat#107860), 40D (cat#60100), 40D-UAS (cat#60101), and UAS-Dronc-RNAi (cat# 100424) were obtained from the Vienna Drosophila Resource Centre. Yw, UAS-myr-RFP, and hs-FLP;;FRT82B were provided by Daniel St. Johnston, hh-Gal4/TM6b was provided by Jean-Paul Vincent. MiWO fly lines (genotype: hs-FLP, UAS-CD8-GFP;; RpS3[Plac92], act>RpS3>Gal4/TM6b) were described in^12^

### Dissection, fixation, and immunofluorescence

All larvae were washed once and then dissected at room temperature in phosphate-buffered saline (PBS). Hemi-larvae were then immediately transferred to pre-chilled PBS on ice and then fixed in 4% formaldehyde for 20 minutes at room temperature on a nutating mixer. Hemi-larvae were subsequently permeabilized for 20 minutes in room temperature 0.25% Triton X-100 in PBS (PBST) followed by a 30-minute blocking step in 4% foetal bovine serum in PBST. Rabbit anti-DCP1 antibody (Cell Signalling, cat#9578S) was diluted in blocking buffer, and primary incubations occurred overnight at 4°C on a rocker. Larvae were then washed three times for at least ten minutes in PBST at room temperature. Secondary antibody and DAPI were diluted in blocking buffer, and hemi-larvae were incubated in secondary antibody for a minimum of 45 minutes at room temperature, followed by 3X additional 10-minute PBST washes and then mounted in VactaShield mounting medium and sealed with a coverslip and nail varnish.

### Image acquisition

All slides were imaged on Leica SP-5 or SP-8 confocal microscopes using a 40x, 1.3 numerical aperture PL apochromatic oil immersion objective with Leica type F fluorescence immersion oil. Wing discs were imaged at 1.4x digital zoom over the entire pouch region, with Z-steps corresponding to 1µm. All images were captured at either 512×512 or 1024×1024 resolution.

### Statistical tests

For univariate statistics, parametric assumptions were evaluated using a Shapiro-Wilks normality test and a Fligner-Kileen homogeneity of variance test. If parametric assumptions were satisfied, a two-tailed student’s t-test was performed along with a Cohen’s d or Hedge’s g effect size. If assumptions were violated, a two-tailed Mann-Whitney U test was performed along with a Cliff’s δ effect size. A pairwise two-tailed Wilcoxon signed-rank test was performed on wing discs with a competing anterior and non-competing posterior compartment, as the datasets were paired and assumptions were violated. Manual quantifications used for validation of the pipeline were made in FIJI 1.53d and statistical tests were run in Graphpad Prism 8.4.3 using the same workflow.

For the multiple logistic regression analysis shown in Figure 6, the dependent variable was, for each *RpS3*^*+/-*^ cell in all wing discs, whether or not the cell was undergoing apoptosis, as determined using combined single cell and foci segmentation in PECAn. Multiple logistic regressions were then performed in PECAn considering a range of non-multi-collinear terms (as determined by a variance inflation factor below 5) within the PECAn data outputs, and we filtered between alternate models by selecting for the lowest Akaike information criterion. Data transformations were considered but were not seen to noticeably improve the model. To correct for multiple comparisons, p-values were adjusted using the False Discovery Rate (FDR) technique. The same optimization was performed for the logistic regression analysis shown in Supplementary Table 3, however the dependent variable was instead the number of apoptotic vs. non-apoptotic cells in the loser clone border for each wing disc.

Source data for all quantifications and replicates is provided in the Supplementary Source Data File.

## Figure Legends

**Supplementary Figure 1:**
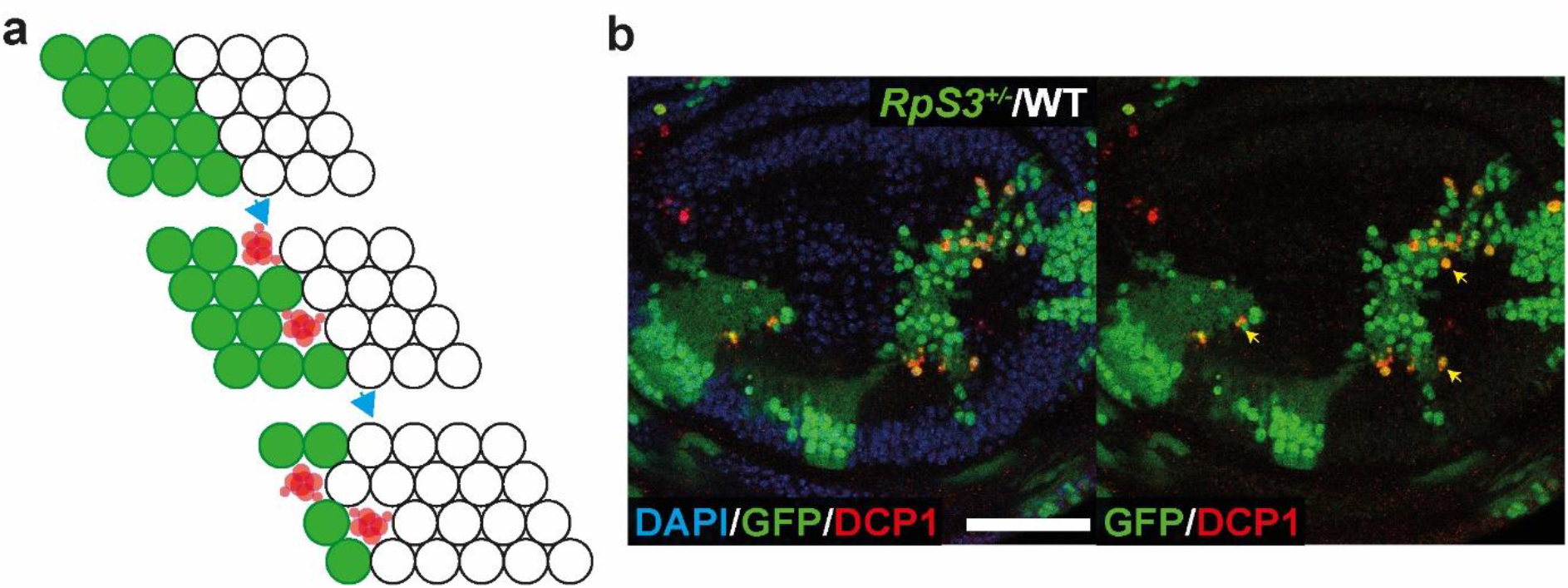
Minute cells are apoptotically eliminated by proximal wildtype cells in mosaic tissues. (**a**) Schematic representation of Minute cell competition. Cells carrying heterozygous mutations in ribosomal protein genes (green) undergo competition when confronted with wild-type cells (white). Minute ‘loser’ cells closest to the wild-type undergo apoptosis (red). Wild-type ‘winner’ cells proliferate and fill the vacated space. Over time, the winner population expands, and the loser population contracts. (**b**) An example of Minute competition in a wandering third instar Drosophila wing disc. Cells heterozygous for *RpS3* (green), a Minute mutation, undergo apoptotic cell death (red, yellow arrowheads) when adjacent to wild-type cells (unlabelled). Scale bar corresponds to 50 µM.

**Supplementary Figure 2:**
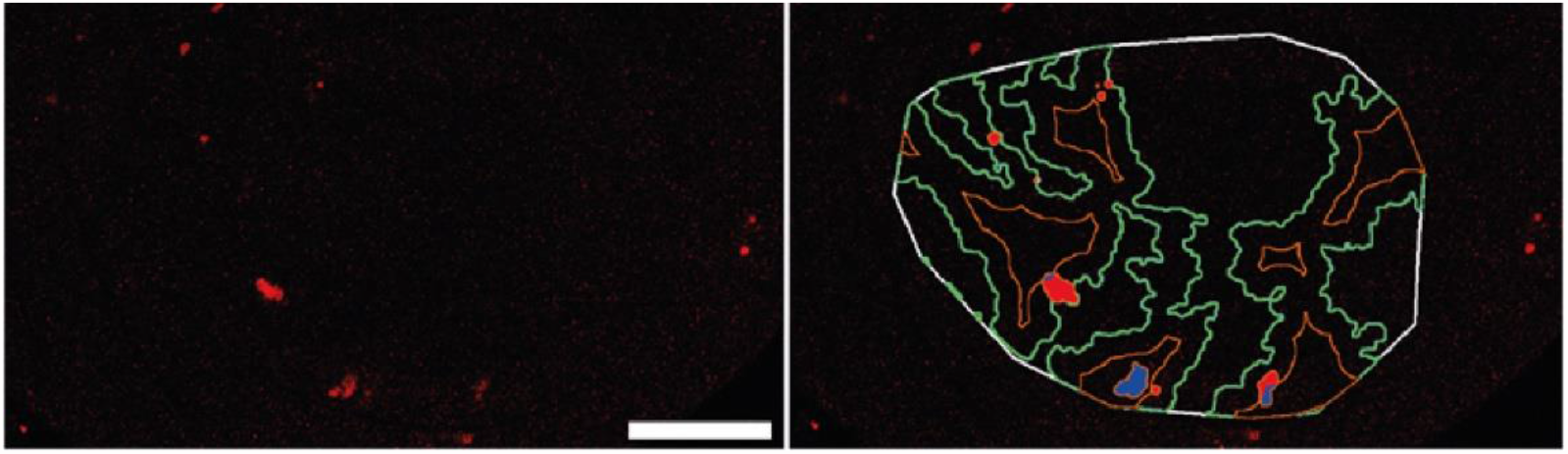
Representative imagej/FIJI PECAn outputs showing cell death segmentation. (Left) The raw cell death channel used for segmentation, along with a 50µm scale bar. This image reflects a MiWO wing disc stained with anti-DCP1 (red). (Right) Overlay of parameters measured by the macro on the raw cell death channel. The white line demarcates the outline of the pouch region. The clone perimeter is shown in green, and the clone border/clone centre boundary is in orange. Cell death signal is automatically assigned as occurring in the border region, shown in red, or as occurring in the clone centre, shown in blue.

**Supplementary Figure 3:**
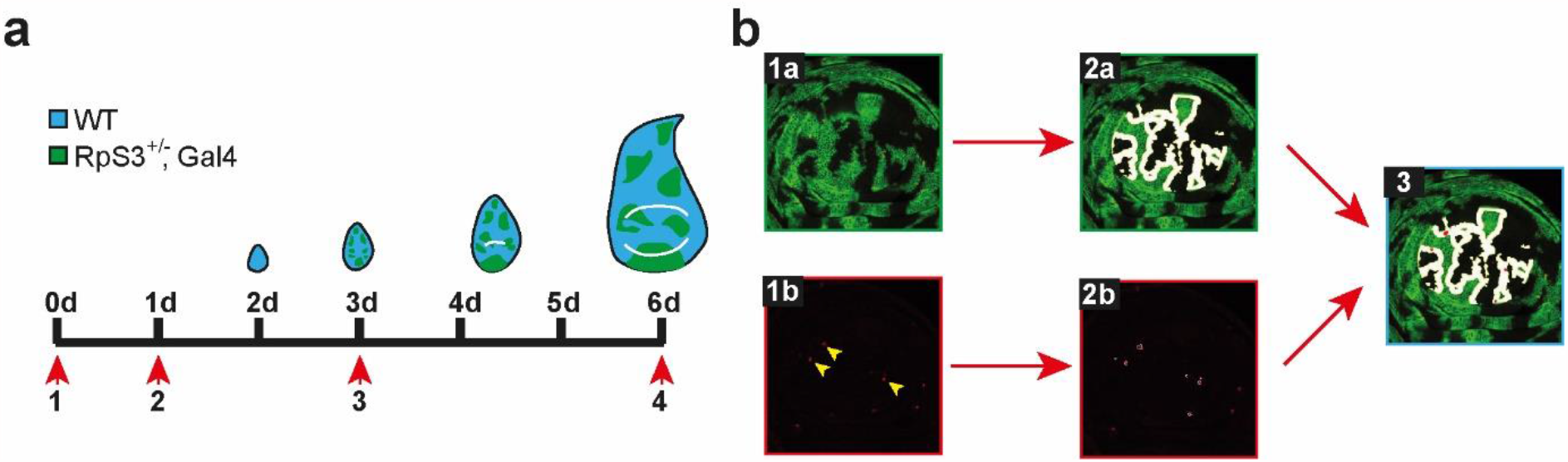
Analysis of Minute competition using the MiWO fly. (**a**) Schematic showing experimental procedure for induction of somatic clones in the study of cell competition using the MiWO tool (Described in ^12^). Although MiWO larvae carry a heterozygous mutation in the *RpS3* ribosomal gene, they hatch as phenotypically wildtype due to the presence of a rescuing transgene driven by an actin promoter, flanked by FRT sites (shown in blue). On day three after egg laying, the larvae are heat-shocked to drive transient expression of the FLP recombinase transgene, which is under control of the hsp70 promoter. The FLP recombinase then drives recombination at the two FRT sites, resulting in the excision of the RpS3 rescuing transgene, as well as a stop codon. The cells are then genotypically Minute, and the actin promoter drives expression of a Gal4 transcription factor, which in turn leads to expression of a UAS-driven GFP (shown in green). Over subsequent days, the cells begin to undergo competitive elimination due to their wildtype winner neighbours. (**b**) Strategy for quantitative analysis of cell competition. In order to assess competition-induced cell death, information is required about both the loser clones (top) and levels of cell death (bottom). First, the clones are segmented. Then they are further segmented automatically into border (the region within two cell diameters of the winners) and clone centre (the region within the clones further than two cell diameters from the winners), where loser cells are shielded from competition. Then, the numbers of dying cells in the border and centre regions are counted. This information is combined to obtain densities of apoptotic events in the clone border versus the clone centre.

